# MMR: A tool for Read Multi-Mapper Resolution

**DOI:** 10.1101/017103

**Authors:** André Kahles, Jonas Behr, Gunnar Rätsch

## Abstract

**Motivation:** Mapping high throughput sequencing data to a reference genome is an essential step for most analysis pipelines aiming at the computational analysis of genome and transcriptome sequencing data. Breaking ties between equally well mapping locations poses a severe problem not only during the alignment phase, but also has significant impact on the results of downstream analyses. We present the multimapper resolution (*MMR*) tool that infers optimal mapping locations from the coverage density of other mapped reads.

**Results:** Filtering alignments with *MMR* can significantly improve the performance of downstream analyses like transcript quantitation and differential testing. We illustrate that the accuracy (Spearman correlation) of transcript quantification increases by 17% when using reads of length 51. In addition, *MMR* decreases the alignment file sizes by more than 50% and this leads to a reduced running time of the quantification tool. Our efficient implementation of the *MMR* algorithm is easily applicable as a post-processing step to existing alignment files in BAM format. Its complexity scales linearly with the number of alignments and requires no further inputs.

**Supplementary Material:** Source code and documentation are available for download at github.com/ratschlab/mmr. Supplementary text and figures, comprehensive testing results and further information can be found at bioweb.me/mmr.

## 1 Introduction

Addressing the increasing need for fast and accurate mapping of high throughput sequencing data to a reference sequence, many different software tools have been developed over the past years, many of which are frequently updated and improved [10, 6, 3, 7]. While numerous challenges have been addressed by the developers, e.g., the consideration of gaps and mismatches or the spliced alignment of RNA-Sequencing data, the problem of ambiguous read mapping still remains unresolved for many of the most popular alignment tools. Depending on factors like read length, alignment sensitivity, and repetitiveness of the target genome, a large fraction of reads aligns uniquely to the target and exactly one mapping location is reported. However, for the remaining, still significantly large, fraction of reads (*≈*10−20%, depending on alignment sensitivity), several possible mapping locations exist. Currently, different strategies are employed to deal with these reads in downstream analyses, most of which have unfavorable side effects: Discarding reads with ambiguous alignments from the alignment result leads to a systematic underestimation of abundance in genomic regions with multi-mapper ambiguities, whereas picking a random alignment or distributing weight across all alignments uniformly does not have a proper biological justification.

Here, we present a simple, yet powerful tool, called the <u>M</u>ulti-<u>M</u>apper <u>R</u>esolution tool (*MMR*), that assigns each read to a unique mapping location in a way that the overall read coverage across the genome is as uniform as possible. *MMR* makes use of the critical fraction of unambiguously aligned reads and iteratively selects the alignments of ambiguously mapping reads in a way the overall coverage becomes more uniform. We show that this strategy has a positive influence on downstream analyses, such as transcript quantification and prediction.

## 2 Approach

### 2.1 Outline of Algorithm

Our approach to resolve ambiguous mapping locations is based on the simple assumption that, besides all existing biases from library preparation and sequencing, the alignment coverage should generally be uniform within a region (RNA-seq or whole exome-seq) or the whole genome (WGS-seq). Based on this assumption, we can evaluate the fit of an alignment of a read to its current mapping location relative to other locations, by assessing the local coverage of the candidate regions. For each read the algorithm jointly evaluates all available alignments with the goal of selecting the alignment/mapping that results in the smoothest overall coverage. At the beginning, for each read one alignment is selected based on either best alignment score, the given input order or random choice. The set of all initially picked alignments as well as alignments of uniquely mapped reads define a global coverage map. Based on this map, we can evaluate the quality of an alignment in its coverage context. To choose the locally optimal alignment for each read, we perform a comparison of all alignments *a* with respect to a loss function ℓ^+^(*a*) of placing *a* relative to not placing it (ℓ^−^(*a*)). In the simplest case the loss function is defined as the empirical variance of the read coverage within window around the alignment (see Suppl. Material). This quantity can be computed efficiently since we keep track of the global coverage map, which is updated when the selected alignment changes. Given the currently selected alignment *a* and an alternative alignment *b*, we update our choice, if the overall loss across the genome would be reduced by choosing the alternative alignment. This is the case when ℓ^−^(*a*) + ℓ^+^(*b*) < ℓ^+^(*a*) + ℓ^−^(*b*). This is repeated for all reads with ambiguous mapping locations. Several iterations over the whole alignment file improve the results. However, the most striking improvements are achieved within the first three iterations and only slight changes can be observed after that. A more detailed description is provided in the Suppl. Methods section.

### 2.2 Paired-end Reads

Handling paired-end reads in our framework is straightforward: Instead of evaluating two individual mapping locations, the same principle is used to compare two pairs of alignments. After generating a list of proper pairs, where feasibility is determined through orientation, chromosome and reasonable proximity, the list is evaluated the same way as the list of possible mapping locations for single-end reads. This approach is easily adaptable to *n*-tuple of mates for *n >* 2.

### 2.3 Limiting Ambiguity

To find a good trade-off between mapping sensitivity and the number of possible mapping locations, we allow to restrict the list of possible mapping locations. This is achieved by thresholding the difference in edit operations between the best hit and any other alignment. For instance, a filter of 0 would only include alignments as possible mapping locations that have as few edit operations as the best mapping.

### 2.4 Implementation

The *MMR* approach is implemented in C++ and its source code is publicly available for download under http://github.com/ratschlab/mmr. Although it has been tested and optimized for Linux based systems, it can be compiled on other platforms. Parsing of alignment files in BAM format requires *samtools* [9]. We also provide a multi-threaded implementation that keeps the coverage information in common memory, requiring no additional memory if multiple threads are used. The single threaded running time depends on the number of possible mapping locations per read but is on average 30−45 seconds per one million alignments per iteration. Thus, running MMR for three iterations on 100 million alignments takes *≈*20 minutes using 10 threads (Intel Xeon E5-2665 CPU).

## 3 Application

As a proof of principle, we tiled the *A. thaliana* genome with overlapping 50-mers and aligned these 50nt reads back to the genome. This resulted in a non-uniform coverage, in particular near repetitive regions (see Fig. 1A; Suppl. Fig. 3). Using *MMR*, we could fully resolve mapping ambiguities, resulting in the expected uniform coverage of 50 almost everywhere.

**Figure 1:**
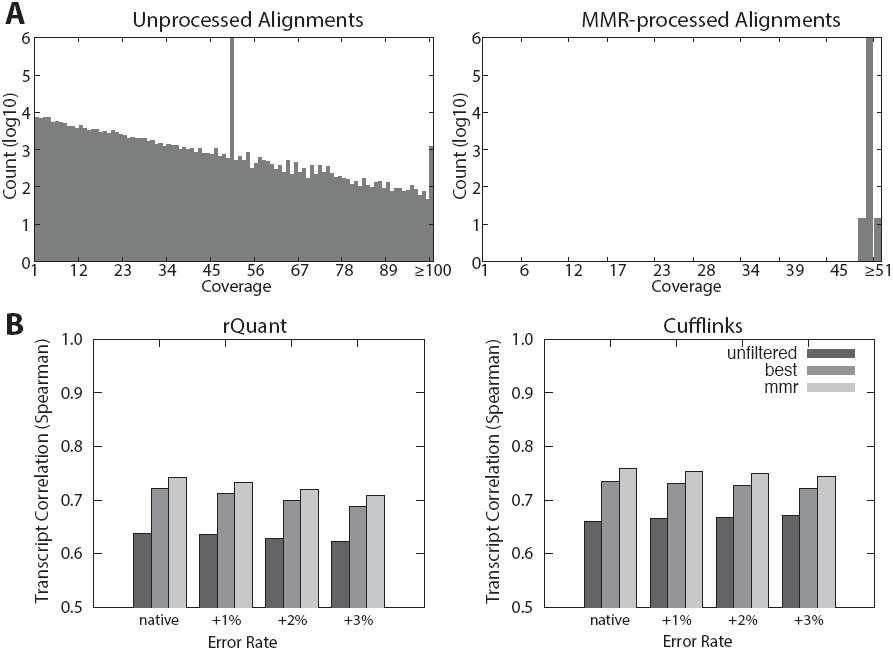
MMR Results on Simulated Read Data: ***A*** *MMR* effect on artificial *A. thaliana* genome tiling data. Distribution of unfiltered coverage values (left) and *MMR*-filtered values (right). ***B*** Quantification results on 3 million simulated 51nt reads aligned with *PALMapper*, using *rQuant* and *Cufflinks*. Accuracy was measured as Spearman correlation to ground truth. Unfiltered read sets are shown in dark, best-hit read sets in medium and *MMR* -filtered in light gray. Native error rate is 2.7%, additional noise levels 1%, 2% & 3%.

We further tested our approach on a set of 3 million artificial RNA-Seq reads that were generated with FluxSimulator [5] based on a subset of 5, 000 genes randomly drawn from the human ENSEMBL annotation. We simulated 51nt and 76nt reads, resulting in an average coverage of 18*×* and 25*×*, respectively. The reads were then mutated using an empirical error model that led to a native error rate of 2.7%. Three levels of random noise (+1%, +2, +3%) were applied in addition. We aligned the reads with *TopHat2* (v2.0.2; [7]) and *PALMapper* (v0.5; [6]), allowing for up to 6 edit operations, with no annotation provided. Further information is provided in the Suppl. Material. To investigate the effect of *MMR* on downstream analyses, we performed transcript quantification using *Cufflinks* [12] *(v1.3) and rQuant* [2] *on the MMR*-filtered alignments, the best alignments only (the alignment ranked highest by aligner) and on completely unfiltered alignments. For *TopHat2* as well as *PALMapper* the quantifications based on the *MMR*-filtered alignments showed a consistently better correlation to the ground truth than both the best-hit and unfiltered alignments sets. The shorter reads of length 51nt (Fig. 1B) showed larger improvements compared to unfiltered (*Cufflinks* : 14.8%, *rQuant* : 16.6%) and best-hit set (*Cufflinks* : 3.2%, *rQuant* : 3.0%) than the longer reads of length 76nt, that showed consistent but smaller improvements (Suppl. Fig. 4 & 5).

## 4 Conclusion

We presented *MMR*, a post-processor for BAM files, resolving ambiguous alignment locations. We showed its easy applicability to the output of different alignment methods and illustrated that *MMR* can greatly improve accuracy of downstream quantification methods. Whereas the improvements seem moderate on a global scale, the effect on single genes can be much larger. Given its lean implementation and the short running time, *MMR* is very well suited for large-scale genome-, exome- and RNA-sequencing efforts. It may also be useful for post-processing alignments in meta-genome projects for improved selection of taxa.

## Acknowledgments

The authors would like to thank Geraldine Jean for fruitful discussions and helpful comments. This work was supported by funding from Max Planck Society, German Research Foundation (RA1894/2-1), Memorial Sloan Kettering Cancer Center and the Lucille Castori Center for Microbes, Inflammation, and Cancer (No. 223316).

## A Methods

### A.1 Description of Algorithm

Following the ideas described in the main manuscript and using the assumption of locally smooth coverage, *MMR* evaluates the whole set of possible alignments for a each given read. The goal is to identify the one alignment for each read that results in the locally smoothest coverage. There are multiple ways to express our prior knowledge in what the coverage should look like. We assume that this prior knowledge is encoded in a global loss function ℓ that computes the global amount of non-smoothness for a set of chosen alignments. In the simplest case, we measure smoothness as the empirical variance of the position-wise coverage in a window of a given length (see (1) below). Other options are discussed in Suppl. Section A.4. The algorithm then minimizes the loss function over all possible alignment choices and chooses the alignment with the smallest overall loss (i.e., greatest “smoothness”). For this, an iterative procedure is applied. Given an input of *k* different alignments for a given read, one alignment is designated as the currently selected one. Depending on user preference this is either an arbitrary alignment or the mapping with the highest alignment quality (see user documentation). The currently selected mapping is then compared to each of the remaining mapping possibilities. For a single comparison of two alignments *a* and *b*, four loss function values are computed: the loss with alignment *a* placed vs not placed and alignment *b* placed vs not placed. Given two possible alignments *a* and *b* to the genomic start locations *pa* and *pb*, respectively, the score 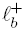 computes the local loss around genomic location *la* if *a* is chosen and 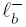 if it is not chosen (i.e., the read is aligned somewhere else); 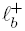 and 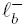 are defined analogously using the alignment *b* to position *pb*.

In the simplest case the loss function is defined as the empirical variance over the genomic coverage of all window positions

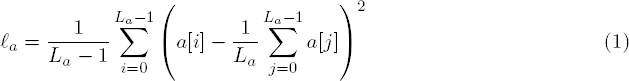

where *L*_a_ is the length of the window around alignment (influenced by option `-w`), *a* [*i*] indicates the read coverage at position *i* relative to the start *pa* of the alignment *a*. If an alignment is present within the window, it influences the coverage and thus the local variance or more generally the loss function. After computing all four values, alignment *a* is chosen if

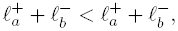

otherwise alignment *b* is chosen. Figure 2 shows a schematic of the *MMR* principle. We discuss other loss functions in Suppl. Section A.4.

**Figure 2:**
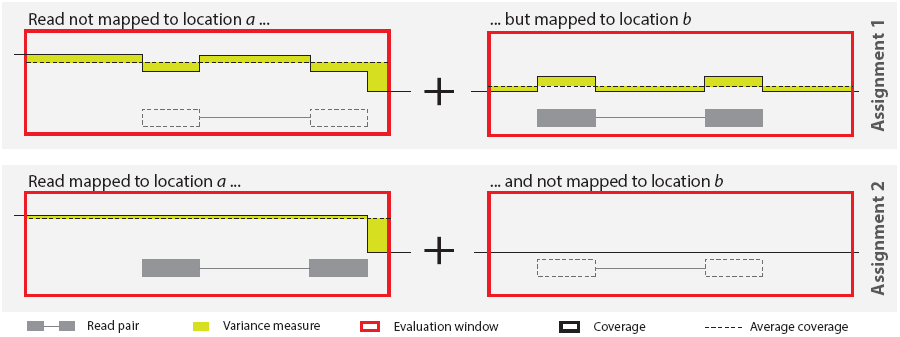
Schematic of the MMR Principle. — Schematic overview of the principle to resolve ambiguous read-mappings. The candidate read-pair in gray has two possible alignments *a* starting at position *p*_*a*_ (left) and *b* starting at position *p*_*b*_ (right). Loss measures (yellow) are computed for both locations, with and without the read-pair. Loss values from the text have following correspondents in the schema: 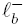 location *a* (top), 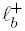 − location *a* (bottom), 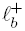 − location *b* (top), 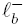 − location *b* (bottom). The evaluation windows are shown in red and the coverage of placed reads as black solid lines.

### A.2 Overlapping Alignment Locations

A major complication arising during the computation of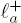, 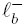, 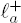 and 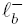is the special case that occurs when the windows of *a* and *b* share common positions. In this situation, two different scenarios can occur:

1. the windows share positions but the alignments do not share positions,
2. the alignments share positions.

As the read is placed at either the one or the other location, in case 1) the computation of 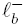 needs to consider coverage contributed by *b* as this will be placed instead of *a* and 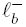 needs to consider coverage contributed by *a*. Case 2) causes a subset of positions that are shared by *a* and *b* to not be altered by the decision. These positions can be masked for analysis and left out in computation, as they contribute to both locations not changing the result.

### A.3 Paired-end Reads

The approach described above can be easily extended to also work for paired-end alignments. In this case, a preprocessing step iterates through all possible valid pairs of alignments of the two mates (see option `-p`). An alignment pair is valid, if the corresponding alignments do not overlap in a conflicting manner. For instance, a conflict would occur, if the first read-mate is aligned into the intronic portion of the second read-mate, if both reads are aligned in the same direction, if the reads align to different chromosomes, or if both alignments have a distance outside of a user-defined maximum range (see option `-i`). After this preprocessing-step, each alignment pair is treated as single alignment possibility *a*_*k*_ and the algorithm above is applied. As the number of possible pairs is quadratic in the number of alignments in the worst case, the number of allowed pairs can be limited by the user (see option `-A`).

### A.4 Loss Functions based on Segment Annotations

For RNA-seq data, one limitation of the straight-forward strategy described so far is that known transcripts are not taken into account. Especially the exon−intron boundaries show steep changes in coverage, but also within exons a change in coverage can often be explained by a mixed signal from several transcript isoforms that superimpose each other. If the transcripts are (approximately) known, this effect can be accounted for during the optimization process. To include structural information into *MMR*, we devised a strategy that takes transcript annotations and quantifications produced by another tool into account. This method can be applied in an iterative scheme. It starts with transcript isoform prediction/quantifications on the alignments using the best hit. Ambiguous alignments can then be re-evaluated based on the intermediate transcript structure and the estimated transcript expression. The improved alignments can then be used to generate improved isoform predictions and quantifications. This can be repeated a fixed number of times or until convergence of the predicted quantifications. *MMR* allows to input transcript annotations together with transcript quantifications (see option `-s`).

To reposition an ambiguous alignment if transcript structures are given, we devised a strategy for an iterative application of *MMR* and *MiTie* [1], a tool for the prediction and quantification of transcript isoforms. If the exon boundaries of all transcript isoforms of a gene are projected to genomic coordinates, the gene can be cut into a set of non-overlapping exonic segments. Thus, each isoform can be built from a subset of these segments. Several isoforms can share the same segment. The expression value of a single segment is the sum of the segment’s expression values over all transcript-isoforms containing that segment (for a more formal description we refer to the publication of MiTie [1]).

The segments as well as corresponding expression estimates provided by *MiTie* or other tools can be used as input for *MMR* (option `-s`). These segments imply a segmentation for the whole genome. Each given segment is associated with a predicted expression value. The empty genomic regions in between any segments are implicitly turned into segments with a predicted coverage of 0. Instead of minimizing the local variance, we now minimize the difference between the observed coverage in an exonic segment with and without the alignment of question and the predicted coverage of the segment.

Again taking the example of the two alignments *a* and *b*, for each alignment we can now identify all genomic segments it overlaps with. Let alignment *a* overlap the *m* genomic segments *g*_*a*,1_, … , *g*_*a,m*_. We can then compute two coverage values for each segment. The value 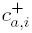 that contains the coverage of segment *g*_*a,i*_ if alignment *a* is mapped to segment *i* and 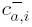 that contains the coverage of the same segment if *a* is mapped to a different location. For each segment we can then compute the difference between observed and predicted coverage, using the expression estimates *e*_*a,i*_ corresponding to the respective genomic segments *g*_*a,i*_ (*i* = 1, *… , m*). Thus, the total loss of alignment *a* is

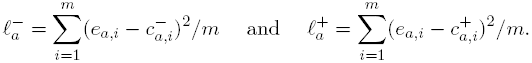

Analogously, for alignment *b* we define the total loss of overlapping genomic segments *g*_*b,i*_ with expression estimates *e*_*b,i*_ (*i* = 1, *… , n*) as

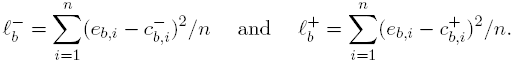

Besides the different calculation of 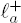, 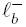, 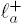 and 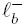, all other steps are identical to the steps described before. Although slightly different adaptations need to be made in order to account for overlapping alignment locations, the same general principles apply.

### A.4.1 Hybrid Strategy

We also implemented a hybrid-strategy that does not rely on predicted expression values for the segments and therefore makes *MMR* much more practical. Instead it uses the minimization of the coverage-variance on a segment-basis and computes the sum over all segments an alignment overlaps, to determine a total variance value. All other other steps are the same as described before (see option `-M`).

### A.4.1 Other Loss Functions

To better account for properties inherent to read-count data, *MMR* can also make use of more general loss functions. For instance, in [1] we used a log-likelihood (loss) function based on a negative binomial distribution. For technical reasons, *MiTie* uses a piece-wise-linear approximation to the log-likelihood loss function. *MMR* supports the specification of an arbitrary piece-wise linear function to score differences between the observed and the expected read coverage (see option `-m`). For further details see user documentation and [1], Suppl. Section K.

## B Results

As described in the main text, we evaluated *MMR* on two different simulated datasets for an assessment of its properties and performance: a) Whole-Genome Sequencing (WGS) in *A. thaliana* and b) RNA-seq in human. All results were obtained using the basic version of *MMR* and without any additional inputs, i.e., no annotation or quantification was used for these experiments.

### B.1 Post-Processing Alignment of Whole-Genome Sequencing for Improved Repeat Handling

For the first dataset we tiled the complete *A. thaliana* TAIR10 reference genome [8] at each position into a set of overlapping 50-mers, thus generating artificial reads from whole genome sequencing, including all low-complexity regions of the genome. In this idealized dataset the coverage at each genomic position (except the 50nt at each end) is exactly 50. We then used PALMapper to realign the first 1, 000, 000 reads back to the *A. thaliana* genome, allowing for up to 5 edit operations, thus generating a high level of additional ambiguity.

### B.2 Post-Processing RNA-seq for Accurate Transcript Quantification

As shown in the histograms of coverage distributions in the main text, *MMR* is able to fully resolve all read-ambiguities in the genomic DNA dataset. In Figure 3, we show an example for a genomic region that shows an uneven coverage before *MMR* filtering and is smoothed after filtering. Notably, the unfiltered alignments showed single genome positions with a coverage exceeding 1, 700 (these are contained in the last bin of the histogram in Fig. 1).

**Figure 3:**
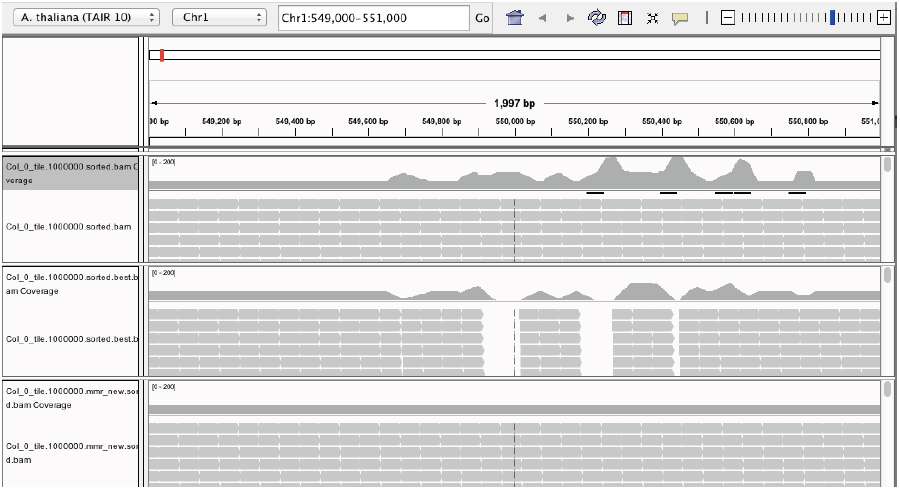
Effect of *MMR* Smoothing — IGV. [11] snapshot of the unfiltered (top), best-hit (middle) and MMR-processed (bottom) alignments. Ambiguous mappings causing unequal distributions could be fully resolved. Missing coverage caused by retaining only the best alignment cannot be observed in the *MMR* processed alignments.

The second evaluation dataset, simulation with FluxSimulator [5], consisted of 3 *·* 10^6^ artificial RNA-Seq reads sampled from 5,000 randomly selected genes of the human ENSEMBL annotation [4]. We simulated two different read-lengths of 51nt and 76nt, resulting in an average coverage of 18*×* and 25*×*, respectively. The reads were mutated as described before and aligned to the hg19 human reference genome using *TopHat* (version 2.0.2 [7]) and PALMapper (version 0.5 [6]) allowing up to 6 edit operations without additional annotation information provided. All other parameters were left at the default.

Based on this dataset, we tested the effect of *MMR* on downstream analyses. For this, we used the unprocessed, the *MMR*-filtered and the best-hit alignment set to perform *in silico* transcript quantification using both *cufflinks* [12] (version 1.3) and rQuant [2], where the best-hit set consisted of those alignments that were ranked highest by the alignment algorithm. Figure 4 shows the quantification results for a read length of 51nt, for all combinations of the aligners and the quantification tools used. Figure 5 shows the respective results for a read length of 76nt.

**Figure 4:**
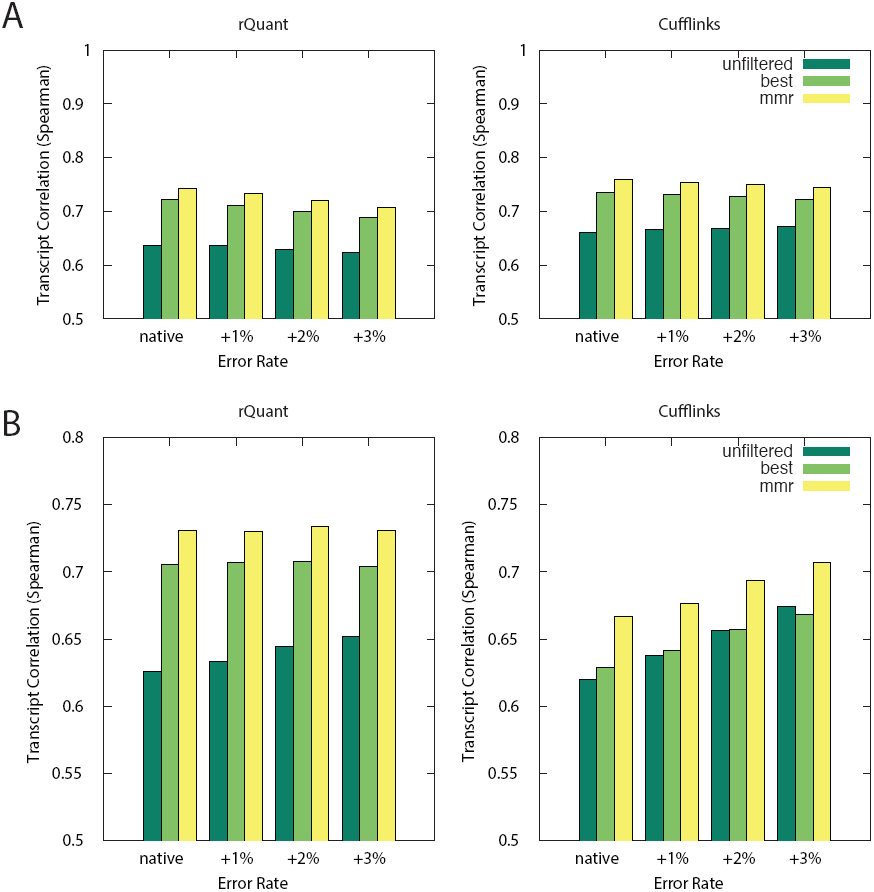
Quantification Results of 51nt reads — A: Accuracy of predicted transcript quantifications by *rQuant* (left) and *cufflinks* (right) measured as rank correlation coefficient (Spearman). Quantification is based on PALMapper alignments. ***B:*** Quantification accuracy as measured before but with quantification based on *TopHat2* alignments.

**Figure 5:**
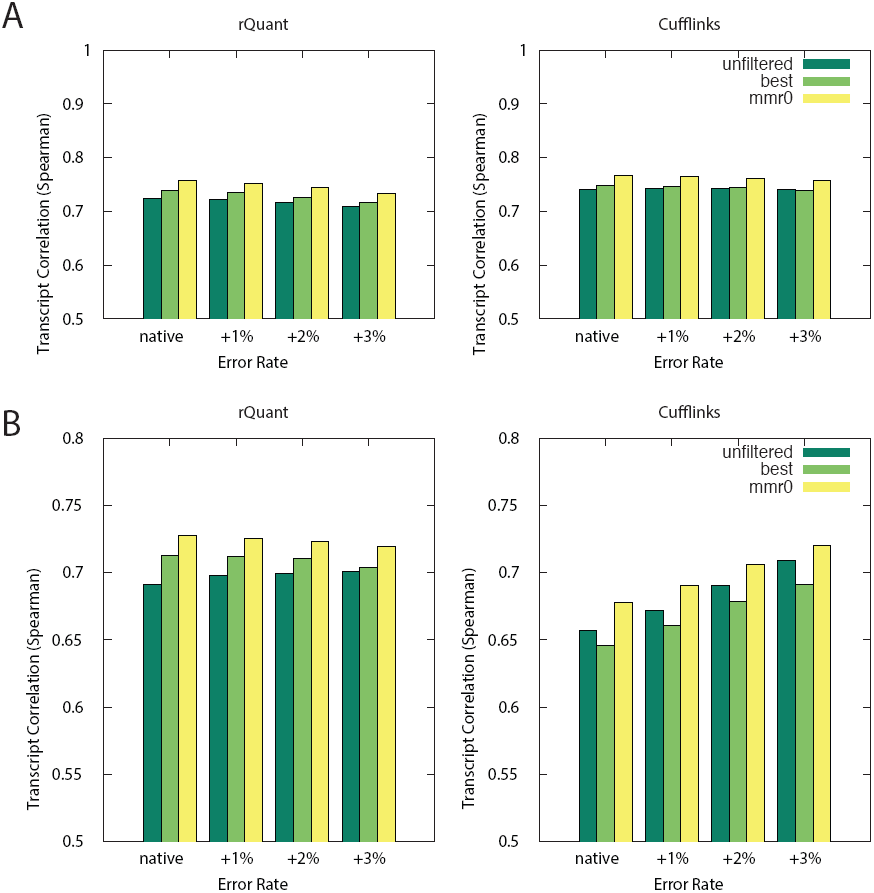
Quantification Results of 76nt reads — A: Accuracy of predicted transcript quantifications by *rQuant* (left) and *cufflinks* (right) measured as rank correlation coefficient (Spearman). Quantification is based on PALMapper alignments. ***B:*** Quantification accuracy as measured before but with quantification based on TopHat2 alignments.

## References

[1] J Behr, A Kahles, Y Zhong, V T Sreedharan, P Drewe, and G Rätsch. MITIE: Simultaneous RNA-Seq-based transcript identification and quantification in multiple samples. Bioinformatics, 29(20):2529–2538, september 2013.

[2] Regina Bohnert, Jonas Behr, and Gunnar Rätsch. Transcript quantification with RNA-Seq data. BMC Bioinformatics, 10(Suppl 13):P5, 2009.

[3] Alexander Dobin, Carrie a Davis, Felix Schlesinger, Jorg Drenkow, Chris Zaleski, Sonali Jha, Philippe Batut, Mark Chaisson, and Thomas R Gingeras. STAR: ultrafast universal RNA-seq aligner. Bioinformatics, 29(1):15–21, january 2013.

[4] Paul Flicek, M Ridwan Amode, Daniel Barrell, Kathryn Beal, Konstantinos Billis, Simon Brent, Denise Carvalho-Silva, Peter Clapham, Guy Coates, Stephen Fitzgerald, Laurent Gil, Carlos García Girón, Leo Gordon, Thibaut Hourlier, Sarah Hunt, Nathan Johnson, Thomas Juettemann, Andreas K Kähäri, Stephen Keenan, Eugene Kulesha, Fergal J Martin, Thomas Maurel, William M McLaren, Daniel N Murphy, Rishi Nag, Bert Overduin, Miguel Pignatelli, Bethan Pritchard, Emily Pritchard, Harpreet S Riat, Magali Ruffier, Daniel Sheppard, Kieron Taylor, Anja Thormann, Stephen J Trevanion, Alessandro Vullo, Steven P Wilder, Mark Wilson, Amonida Zadissa, Bronwen L Aken, Ewan Birney, Fiona Cunningham, Jennifer Harrow, Javier Herrero, Tim J P Hubbard, Rhoda Kinsella, Matthieu Muffato, Anne Parker, Giulietta Spudich, Andy Yates, Daniel R Zerbino, and Stephen M J Searle. Ensembl 2014. Nucleic Acids Research, 42(Database issue):D749–755, 2014.

[5] Thasso Griebel, Benedikt Zacher, Paolo Ribeca, Emanuele Raineri, Vincent Lacroix, Roderic Guigó, and Michael Sammeth. Modelling and simulating generic RNA-Seq experiments with the flux simulator. Nucleic Acids Research, 40(20):10073–10083, september 2012.

[6] G Jean, A Kahles, V T Sreedharan, F De Bona, and G Rätsch. RNA-Seq read alignments with PALMapper. Current Protocols in Bioinformatics, Chapter 11(December):Unit 11.6, December 2010.

[7] Daehwan Kim, Geo Pertea, Cole Trapnell, Harold Pimentel, Ryan Kelley, and Steven L Salzberg. TopHat2: accurate alignment of transcriptomes in the presence of insertions, deletions and gene fusions. Genome Biology, 14(4):R36, April 2013.

[8] Philippe Lamesch, Tanya Z Berardini, Donghui Li, David Swarbreck, Christopher Wilks, Rajkumar Sasidharan, Robert Muller, Kate Dreher, Debbie L Alexander, Margarita Garcia-Hernandez, Athikkattuvalasu S Karthikeyan, Cynthia H Lee, William D Nelson, Larry Ploetz, Shanker Singh, April Wensel, and Eva Huala. The Arabidopsis Information Resource (TAIR): improved gene annotation and new tools. Nucleic Acids Research, 40(Database issue):D1202–1210, January 2012.

[9] H Li, B Handsaker, A Wysoker, and T Fennell. The sequence alignment/map format and SAMtools. Bioinformatics, 25(16):2078–2079, 2009.

[10] Heng Li and Richard Durbin. Fast and accurate short read alignment with Burrows-Wheeler transform. Bioinformatics, 25(14):1754–1760, july 2009.

[11] James T Robinson, Helga Thorvaldsdóttir, Wendy Winckler, Mitchell Guttman, Eric S Lander, Gad Getz, and Jill P Mesirov. Integrative genomics viewer. Nature Biotechnology, 29(1):24–26, 2011.

[12] Cole Trapnell, Brian A Williams, Geo Pertea, Ali Mortazavi, Gordon Kwan, Marijke J van Baren, Steven L Salzberg, Barbara J Wold, and Lior Pachter. Transcript assembly and quantification by RNA-Seq reveals unannotated transcripts and isoform switching during cell differentiation. Nature Biotechnology, 28(5):511–515, 2010.

